# Subtype-specific Patterns of Evolution and Clinically Relevant Genomic Instability in Wilms Tumour

**DOI:** 10.1101/2024.09.04.610994

**Authors:** George D. Cresswell, Tasnim Chagtai, Reem Al-Saadi, Taryn D. Treger, Gaganjit Madhan, Borbala Mifsud, Gordan Vujanic, Richard D. Williams, Nicholas M. Luscombe, Kathy Pritchard-Jones, William Mifsud

**Affiliations:** The Francis Crick Institute, London NW1 1AT, United Kingdom.; UCL Great Ormond Street Institute of Child Health, London WC1N 1EH, United Kingdom.; Great Ormond Street Hospital for Children NHS Trust, Histopathology Department, Great Ormond Street, London WC1N 3JH, United Kingdom.; College of Health and Life Sciences, Hamad bin Khalifa University Doha, Education City, Qatar.; Department of Pathology, Sidra Medicine, Al Luqta Street, Doha, Qatar.; Weill Cornell Medicine Qatar, Doha, Qatar.; NIHR Imperial BRC Genomics Facility, Faculty of Medicine, Imperial College London, United Kingdom.; UCL Genetics Institute, University College London, London WC1E 6BT, United Kingdom.; Okinawa Institute of Science & Technology Graduate University, Okinawa 904-0495, Japan.

**Keywords:** Cancer evolution, genomic instability, paediatric cancer

## Abstract

**Background:** Understanding cancer evolution is fundamental to predicting cancer progression. However, the evolution of paediatric cancers is still under-researched. Large cohorts of patients are required to determine consistent evolutionary trajectories that shed light on key steps in cancer development and reveal underlying biology, especially in rare cancers. Additionally, well annotated clinical data is necessary for determining if evolutionary biomarkers are predictive of patient outcome.

**Methods:** We performed detailed evolutionary analysis of 64 paediatric kidney cancers, including 60 Wilms tumours (WT), using DNA microarrays and, in a subset of 30 patients, a WT-specific targeted sequencing assay, to detect copy number alterations (CNA) and mutations, respectively. By analysing multiple tissue samples in the majority of cases we could detect mutation heterogeneity in each tumour. We reconstructed clones across the cohort and described their phylogenetic histories, in addition to detecting mirrored subclonal allelic imbalance.

**Results:** Our results highlight pervasive evidence of parallel evolution in WTs affecting CNAs, and *CTNNB1* and *TP53* mutations. Furthermore, we demonstrate that stromal-type WTs often evolve from a consistent series of events (*WT1* mutation, 11p uniparental disomy and *CTNNB1* mutation) and we suggest that 19q uniparental disomy is an important ancestral event in both epithelial and diffuse anaplastic WTs. Finally, we propose the total number of evolutionary CNA events as a prognostic biomarker in WTs for event-free survival, particularly in high-risk WT.

**Conclusions:** Overall, this study sheds light on the evolution of the most common paediatric kidney cancer and links evolutionary analysis to fundamental clinical and biological questions in a large cohort of WTs. We conclude that histological subtypes of WT are often defined by consistent evolutionary sequences. We present evidence that a key marker of evolvability, namely CNA diversity measured phylogenetically across multiple tumour sites, is prognostic of patient outcome and should be considered for clinical use to detect the most aggressive blastemal and diffuse anaplastic type WTs.

## 1 Background

Cancer cell populations are subject to the forces of evolution due, in part, to their ability to randomly acquire heritable mutation events that may alter the phenotype of the cell during cell division ^1^. Populations of cancer cells are consequentially genetically diverse ^2^. As in species evolution, the genetic composition of a tumour population is informative of the evolutionary history of a cancer and provides a context for understanding how genetic alterations arose over time and the selection pressures that influenced them ^3^. Research over the previous decade, catalysed by the advent of next generation sequencing, has spurred the deconvolution of adult cancer cell populations and the inference of tumour evolution namely through bulk sample analysis ^2,4^, multiregion assays ^5,6^ and single cell sequencing ^7,8^.

Evolutionary processes in paediatric tumours are less well-understood, due to a combination of factors including disease rarity and relatively low mutational burden ^9^. Nevertheless, we ^10^ and others ^11–15^ have previously shown that paediatric cancer populations are genetically heterogeneous and subject to evolutionary processes; and that these findings are best supported by studies in which multiple samples from a tumour are analysed. Whilst detecting point mutations and small insertions and deletions in driver genes is vital to understanding paediatric cancer, the paucity of point mutations makes it more important to reliably identify copy number aberrations (CNA) and loss of heterozygosity (LOH) events, especially those that are only present within a subset of cells in a tumour sample, in order to understand the evolution of paediatric tumours. Furthermore, it is particularly important yet challenging to study relatively large cohorts of cases with a given childhood tumour diagnosis, especially when the diagnosis is not entirely dependent on a pathognomonic molecular abnormality. In order to address these challenges, we developed methods that allowed us to improve the detection of CNA and LOH events in tumour subpopulations, and used these to study a set of sixty-four paediatric kidney cancer cases (206 tumour samples) of which the majority were multi-sampled (83%). In a subset of thirty cases we also obtained information about point mutations and small insertions and deletions by massively parallel targeted exon sequencing (TES). Specifically, we characterised CNA and LOH events, including subclonal events, in each of the multiple tumour samples, and determined all unique observable clone states across each tumour. We then constructed phylogenetic trees for each tumour. Our cohort included rich clinical annotation, which allowed us to assess the relationships between CNA/LOH burden and types, evolutionary patterns, and tumour histology and clinical outcome.

## 2 Methods

### 2.1 Sample collection

We obtained multiple tumour samples from WT nephrectomy/nephron-sparing surgery specimens from patients enrolled on the Improving Population Outcomes for Renal Tumours of Childhood (IMPORT) study or whose parents had consented for additional tissue to be used in research as part of the UK Children’s Cancer and Leukaemia Group (CCLG) tissue bank. In a subset of cases, we also studied material obtained at diagnostic needle core biopsy. In addition, in one case, tissue from a relapse biopsy was also used. The research was approved by a national research ethics committee (12/LO/0101). Patients received preoperative chemotherapy as per the IMPORT study protocol.

WTs were classified for diagnostic purposes as previously described ^16^. Each research tissue sample was divided into two pieces. One piece was formalin-fixed and paraffin-embedded and from this at least one histological section was prepared, stained with haematoxylin and eosin (H&E stain) and reviewed by one pathologist (Dr. William Mifsud). Standard histological methods, as per standard operating procedures in place at the Department of Histopathology at Great Ormond Street Hospital, were used. The matching piece was flash-frozen in liquid nitrogen. DNA was extracted from frozen tissue samples using standard techniques, and from adjacent non-tumorous kidney tissue and/or peripheral blood lymphocytes where they were available. Cases BI-1, BI-2, BI-4, DA-1, MX-1 and MX-3 were previously presented using an alternative SNP array, without WT-TES sequencing and with less advanced analysis ^10^.

### 2.2 Copy number detection using Illumina^®^ SNP arrays

Illumina^®^ CoreExome-24 SNP arrays (~500,000 probes) were hybridised with 250ng DNA per sample according to the manufacturer’s instructions. Log R ratio (log_2_[observed intensity/reference intensity], LRR) and B-allele frequency (BAF) were calculated using the Illumina^®^ GenomeStudio software for each array using default settings. SNP array data in these array platforms was generated by UCL Genomics.

### 2.3 Targeted NGS of tumour samples using a WT specific panel

In total 181 genes (0.881 Mb) were pulled down for sequencing analysis (Table S1) designed using the Agilent^®^ SureSelect SureDesign suite. These genes were chosen based on known WT genes from the literature ^17^. Maximum tiling density was used to pull down the exons of this gene list (5X).

Library preparation, targeted capture and sequencing was carried out at the UCL Pathogen Genomics Unit. NGS was performed on an Illumina^®^ NextSeq machine. For each sample paired-end reads of 151 base pairs were generated. Sequencing data was provided as FASTQ files.

### 2.4 Copy number quantification

LRR genomic waves ^18^ were detected in normal tissue samples and corrected from all arrays as described previously ^10^. Only autosomes were analysed as male X chromosomes do not possess heterozygous SNPs. LRRs were normalised to the median and outliers were smoothed using CGHcall ^19^. Heterozygous probes were identified by selecting probes in a matched normal sample with BAF values between 0.3 – 0.7 or were manually selected in cases without a good quality matched normal sample. The heterozygous probes BAF values are transformed to mirrored BAF (mBAF). The mBAF were then segmented and transformed to a major allele frequency distribution. The code for this transformation is available as an R package (www.github.com/georgecresswell/MiMMAl). The median BAF of the major allele distribution and mean LRR was used as input to ASCAT^20^ to determine the purity and ploidy of the sample and copy number cancer cell fractions were determined using a modified version of the Battenberg algorithm ^4^ (Supplementary Methods).

### 2.5 Identifying Mirrored Subclonal Allelic Imbalance

K-means clustering was performed in R (k = 2) on the BAF of each genomic region of CNA breakpoints across multiple samples. Allelic imbalance was called when the mean BAF of two clusters in a sample had a difference *>*0.1 and if two samples showed allelic imbalances in opposite directions mirrored subclonal allelic imbalance (MSAI) was called.

### 2.6 Mutation calling

FASTQ files were trimmed using cutadapt ^21^. Reads were aligned to the human genome (GRCh37.p13) using bowtie2^22^. Bowtie2 was run using the settings --sensitive --score-min L,0,-0.12. Samtools was used to output mapped reads into sambamba (www.github.com/lomereiter/sambamba) which was used to generate a sorted bam file ^23^. Duplicated reads were then marked using picard tools (www.broadinstitute.github.io/picard). Point mutations in bait capture regions were called using MuTect version 1.1.7^24^. Calls filtered by ‘nearby gap events’ and ‘alt allele in normal’ alone were passed. Indels were called using MuTect2 in the Genome Analysis Tool Kit version 3.6^25^. MuTect2 was used to analyse the targeted pull down regions of the genes *CTNNB1*, *WT1*, *AMER1*, *DICER1*, *DGCR8*, *TP53*, *DROSHA* and *MLLT1*. Calls filtered by ‘str contraction’ and ‘alt allele in normal’ alone were passed. The dbsnp138 release as reference for the option --dbsnp. Mutations were kept if there were minimum 10 reads reporting the mutation and the variant allele frequency (VAF) in the tumour sample was 10x greater than in the normal sample. The VAF of each mutation of a patient was then reported for all regions. Mutations were annotated using Oncotator ^26^.

### 2.7 Defining clones and phylogenetic analysis

Clones in each individual sample were defined using Gaussian density smoothing and in segments with a mixture of states we assumed that the state closer to diploid occurred earlier in the phylogenetic history of the tumour (Supplementary Methods). Clones identified within each sample were then interpreted as being nested in all phylogenetic trees apart from manually curated exceptions with strong evidence of clones within a sample belonging to separate lineages. Clones were then used as input for MEDICC ^27^ to generate phylogenetic trees based on copy number events. All trees and calls were manually assessed and curated based on the reinterpretation of CNA events in the phylogenetic trees not considered by MEDICC such as separate CNA events with overlapping intervals and MSAI (Supplementary Methods).

### 2.8 Receiver operating characteristic curve and survival analysis

Follow up was available for 53/64 cases. CNA events and their subclonality (percentage non-clonal events) was counted for each CNA phylogenetic tree split by laterality. In bilateral cases the tumour with the largest number of events or subclonality was considered representative of the case. Tumours with a subclonality of 100% (independent tumours) were removed from the predictive assessment of subclonality.

Receiver operating characteristic (ROC) curve analysis was performed using the pROC R package ^28^ using the event-free survival status as the response and the total number of CNA events or the percentage of subclonal CNAs as the predictor. The area under the curve was calculated using the auc() function in the pROC package. The threshold to split the cases by the predictor was calculated by measuring the part of the ROC curve furthest from the function that represents a random predictor (*y* = *−x* + 1). The Kaplan-Meier curves and log-rank test were calculated and performed using the survival R package using the survdiff() and survfit() functions ^29^.

## Supplementary Methods

### Segmentation of SNP arrays

Initially the mBAF for each array is then segmented using piece-wise constant fitting (kmin = 3, gamma = 4.72) on whole chromosomes and these segments are used to determine the ‘sample *sd*’ (see ‘Mixture modelling of the major allele’). Segmentation of the mBAF is then also performed on separate chromosomal arms. Firstly the distance of each mBAF value to the mean mBAF of its segment is measured for each probe and summed for each chromosome. The distance for the two versions are compared and if the distance of the centromere split segmentation is greater the whole chromosome segmentation plus 0.58, the whole chromosome segmentation is retained. If the centromere split segmentation is used, the distribution of BAFs in the segments which lie immediately adjacent of the centromere are tested to be significantly different using the Kolmogorov-Smirnov test. If the distributions are significantly different (*p<*0.05) the centromere split is retained as the final segmentation output, if not, the whole chromosome segmentation is used.

### Mixture modelling of the major allele

A two component Gaussian mixture model is fitted to the BAFs of each segment using expectation maximisation (EM) and the normalmixEM() function in the mixtools R package version 1.0.4^30^. Means are constrained to be equidistant from 0.5 and the standard deviation is constrained to be equal for both components. The starting value for the component means is the mean mBAF and 1 – mean mBAF. The value of *sd* converged upon for each segment is recorded across the array. The values of *sd* are smoothed using kernel density smoothing using the basic R function density() with default settings for bandwidth (BW) calculation. The *sd* of the highest peak of this density is taken as the representative *sd* of the array and a range of ±0.33 either side of the ‘sample *sd*’ is chosen for parameter testing.

Then, for each segment the BAF distribution is tested for a normal BAF distribution (mean = 0.5), using the expected normal distribution given the ‘sample *sd*’ using a Kolmogorov-Smirnov test. If the test shows significant difference (*p<*0.05) then the segment proceeds to be modelled as two mixtures using two parameter grid searches. A log-likelihood is calculated for combinations of distance from 0.5 for the means of the distributions (values for distance from 0.5, i.e. distribution means, are used in the range of 0 – 0.5 in intervals of 0.01) and *sd* of those distributions (20 equal intervals in the *sd* range) and the highest log-likelihood is chosen for the given segment. Conducting a grid search ensures all extreme values are tested (such as the means of both distributions being equal, i.e. distance = 0). Following this first ‘global’ grid search, a ‘local’ grid search is implemented in a more refined range of values surrounding the chosen solution to the global grid search. The ‘local’ search uses the values in the grid immediately neighbouring the ‘global’ solution using 10 intervals between the *sd* values and 20 intervals for the values of the means of the distributions. The combination of parameters producing the largest log-likelihood in the ‘local’ search is taken as the final solution for determining the means and *sd* of the major/minor allele distributions in the segment.

Once two distributions have been fitted to the BAFs in the segment a ‘minor allele ratio’ is calculated using the densities of these two normal distributions given the chosen means and *sd*. The distribution with a mean *<*0.5 is considered to be the minor allele distribution and the distribution with a mean *>*0.5 is defined as the major allele distribution. The ‘minor allele ratio’ is calculated for a given BAF value as the density of minor allele distribution divided by the sum of the densities of the minor and major allele densities. This ratio is then considered to be representative of a probability that a SNP with a given BAF belongs to the minor allele distribution. Each SNP in the segment is subsequently randomly mirrored at 0.5 (1 – BAF) according to the probability that it is in the minor allele distribution. This then transforms the BAFs across the segments into a representation of the major allele distribution.

### Copy number quantification

The median BAF and the mean LRR of each segment are then inputted into the ASCAT workflow to determine the purity, *ρ* and ploidy, *ψ*, of the sample as well as the array gamma (a multiplier that represents the compression effect in the SNP array as described by van Loo and colleagues), *γ*, using the ASCAT equations that model LRR and BAF^20^. In short this involves calculating major and minor allele copy number states across a grid search of *ρ* (0.5 – 1) and *ψ* (1.6 – 3.4) values. These copy numbers are then measured genome-wide for distance from integer states. We altered this workflow by also performing this across a range of *γ* values and selecting the grid search with closest solution to an integer state, in order to also select for the most appropriate *γ* (0.25 – 0.5). The grid is then searched for local optima and the combination of *ρ* and *ψ* is reported for each local minima. We then selected for the solution within a range of accepted *ψ* and *ρ* values that produced the highest *ρ*. These parameters were adjusted in particular cases which showed likely tetraploidy. In general diploid solutions were taken unless segments produced combinations of values impossible in a diploid state (e.g. allelic balance in the BAF but a loss-like negative LRR value – a [1+1], diploid, state in a [2+2], tetraploid, genome). The chosen *ρ* (purity), *ψ* (ploidy) and *γ* are then used to model copy number state mixing.

The modelling of mixed copy number states was performed according to the method of Nik-Zainal and colleagues which is an extension of the ASCAT equations known as ‘Battenberg’ ^4^. We choose to ensure the first two state solution tested by this method was between a normal state [1+1] and the closest aberrant state, prior to testing the remaining combinations as determined by the Battenberg methodology. Therefore for each subclonal segment, a series of two state copy number combinations are reported as cancer cell fraction mixtures, with the first combination of [1+1] and the closest aberrant state taken as the combination with first priority.

### Exceptions for higher than diploid ploidy states

Two cases clearly harboured tetraploid genomes, these were MX-11 and DA-8. Here a larger range of *ψ* was tested in order to allow for a tetraploid solution (1.6 – 5.5). To settle on a tetraploid solution in the R2 sample of MX-11, the range of tested *ρ* values was also expanded (0.2 – 1) in this case and R2 of MX-11 reached a tetraploid solution with a purity of 49%. Also for both solutions in MX-11 the best fitting local minima solution was taken, not the solution with the highest *ρ*.

Owing to the fact that the background in tetraploid cases is no longer a [1+1] state, the two state mixture of [1+1] and an aberrant state is no longer taken as the first option and the the ordering of states is taken entirely according to the Battenberg methodology.

### Clustering copy number alternations to identify subclones

Once the cancer cell fraction of each copy number change has been calculated, each individual array was then analysed to identify clusters of CNAs that may have formed from the same clonal expansion and therefore represent the same clone. We performed this using kernel density estimation. Firstly, segments with a mixed copy number state of two aberrations are assumed to have acquired the aberration with a copy number state genetically closer to [1+1] earlier and this state is considered to have been previously clonal and considered to have 100% purity.

Next Gaussian density smoothing is performed using the density() function in R (bandwidth = 0.08). Troughs in the kernel density estimation are identified and used as boundaries between groups of CNAs. CNAs within these boundaries are then grouped together and considered to be part of the same ‘clone’ and the peak density of each of these clones is taken as the cancer cell fraction of the clone.

As we forced Battenberg to estimate combinations of a normal state [1+1] and the nearest aberrant state as the first candidate for a mixture of copy number states in a non-clonal segment, cancer cell fractions for the aberrant state that exceed 100% were generated in a subset of segments. If these segments produced a ‘clone’ with a cancer cell fraction greater than 110% the ‘second’ option was considered for all CNAs clustered within it (first Battenberg option that is not a combination including [1+1]). As this will be a mixture of two ‘aberrant’ states, the state closest to [1+1] is considered to have once been clonal and the cancer cell fraction is then considered to be 100%. The identification of clones using kernel density smoothing is then repeated and all clone cancer cell fractions will be *<*110%.

If, in a segment with two aberrant states, the state furthest from [1+1] has a very high cancer cell fraction it is possible for it to also be included in the clone with the highest cancer cell fraction. As the other state is also considered part of the clonal peak this would lead to two states existing simultaneously in the clonal peak for the same segment. Therefore in this situation the state furthest from [1+1] is used in the clonal peak.

### Inferring phylogenetic relationships in multiple tumour regions with multiple clones

Once clones have been identified within each tumour array across multiple samples, phylogeny is inferred. For an initial interpretation of phylogeny the clones within each sample are considered to be ‘nested’ even if they can, through virtue of the ‘pigeon hole principle’, be on separate branches in cases with 3 or more subclones in a single sample. However, the ‘pigeon hole principle’ was considered later in the manual inspection of the phylogenetic tree. The breakpoints of the CNAs across all inferred clones were then smoothed and the genome was split into intervals across with to compare a single CNA state using CGHregions (*c* = 0) ^31^. Two separate inputs were then used for running MEDICC ^27^, all inferred clones and only the ‘clonal’ profiles (CNAs in each sample with the highest cancer cell fraction).

This was performed as MEDICC treats each copy number profile used as input entirely separate and was developed for interpreting copy number profiles taken as averages from separate samples. To this effect it is unable to understand the predetermined phylogenetic relationship between two clones identified in a single sample, i.e. that the clone with the higher cancer cell fraction is assumed to have arisen prior to the clone with the lower cancer cell fraction and that this clone is considered to be ‘nested’ inside the higher cancer cell fraction clone. Therefore each consensus phylogenetic tree produced required manual assessment of both approaches, using all identified CNA profiles (all ‘subclones’) and the phylogenetic tree produced by only the ‘clonal’ CNA profiles of each sample (the cluster with the highest cancer cell fraction) as reference. The MEDICC phylogenetic tree could then be assessed for misordering of nested clones when all CNA profiles were used as input and the MEDICC tree produced using only the CNA profiles generated from the highest cancer cell fraction cluster per sample could be interpreted as a possible ‘backbone’ of clonal phylogenetic changes and the nested ‘subclonal’ profile could be mapped on as extension of this backbone.

However, further considerations were also taken when manually assessing the output of MEDICC. These included:

- CNAs affecting largely the same region with clearly different breakpoint boundaries were mostly interpreted as being separate events affecting the same region as opposed to the shortening or lengthening of a pre-existing CNA.
- Breakpoints that were very close but inaccurately identified due to tumour sample impurity were manually defined to be the same if another sample clearly showed a purer version of the same CNA when inspecting profiles.
- Some CNA mixture combinations were re-interpreted if such a change was supported by another sample. Often if the CNA mixture was considered to change the total copy number state but did not alter LRR, it was re-interpreted as a mixture of 2+0 and 1+1, this was often the case for the mis-identification of 11p LOH as a mixture of 2+1 and 1+1 or 2+0 and 2+1.
- If CNAs across a chromosome overlapped they were considered for the possibility of being a mixture of two overlapping CNA states in separate clones.
- The detection of MSAI (a CNA affecting a different allele) was used to separate CNAs as being separate events, even if the breakpoint boundary was equal.

Some sets of CNAs were too challenging to manually assess, such as regions of possible chromothripsis and MEDICC was used to interpret these changes. In some cases WGD makes interpreting CNA evolution difficult (MX-11 and DA-8), and for these cases the interpretation of CNA evolution was carried out by using the results of MEDICC using only the highest cancer cell fraction clones in each sample and then mapping the additional within-sample clones on to the tree.

## 3 Results

To explore copy number evolution in WT we studied a large cohort of patients with the entire range of WT types. In total, we analysed sixty-four patients with paediatric kidney tumours, including sixty with WT; the remaining four patients presented with histologically proven diffuse hyperplastic perilobar nephroblastomatosis (DHPN), multifocal perilobar nephrogenic rests (without WT or DHPN), metanephric adenofibroma and renal cell carcinoma, respectively. Thirteen cases of WT presented as bilateral WT and tumour masses from both kidneys were sampled in more than half of these patients (7/13). In the other six patients, only one side was subjected to nephrectomy/nephron-sparing surgery, on the basis of a radiological diagnosis of contralateral nephrogenic rest(s) rather than WT. Multiple samples were assayed for the majority of WT patients (~83%, 53/64) and the largest number samples taken from a single patient was seven (Table 1). Considering bilateral tumours separately, the most frequent histological types of WT we analysed were mixed, stromal and diffuse anaplastic (Table 2). All samples were assessed for copy number changes using Illumina CoreExome-24 SNP arrays, a subset of patients (n=30) were assessed for mutational heterogeneity using a specifically designed WT targeted panel (WT-TES, Table S1).

**Table 1:** A summary of the number of tumour samples per case and per kidney (treating each tumour in bilateral cases as separate unilateral tumours)

| Number of Samples | By Patient | By Kidney |
| --- | --- | --- |
| 1 | 11 | 13 |
| 2 | 19 | 25 |
| 3 | 8 | 10 |
| 4 | 7 | 9 |
| 5 | 9 | 6 |
| 6 | 7 | 6 |
| 7 | 3 | 2 |

**Table 2:** A summary of the tumour types in the cohort of 64 paediatric kidney cancer cases. The majority of these cases were WT (~94%) and the individual types of WT are also noted, firstly for only the unilateral tumours (Uni. tumours), and secondly including the types of each individual side from the 15 bilateral cases (Uni & Bi.). Subtypes defined by SIOP classification after neoadjuvant chemotherapy ^16^.

| Tumour Type | Type | Uni. tumours | Uni. & Bi. |
| --- | --- | --- | --- |
| Wilms tumour | Blastemal | 6 | 6 |
|  | Diffuse Anaplastic | 9 | 9 |
|  | Epithelial | 5 | 9 |
|  | Focal Anaplastic | 1 | 1 |
|  | Mixed | 12 | 21 |
|  | Necrotic | 2 | 2 |
|  | Regressive | 5 | 6 |
|  | Stromal | 7 | 13 |
| DHPN | - | 0 | 1 |
| Multifocal Perilobar Nephrogenic Rests | - | 0 | 1 |
| Metanephric Adenofibroma | - | 1 | 1 |
| Renal Cell Carcinoma | - | 1 | 1 |

To interpret copy number evolution in WT we conducted phylogenetic analysis in all patients by assessing copy number alteration (CNA) in multiple samples and assigning each CNA a cancer cell fraction (CCF). CNAs with similar CCF were clustered together and inferred to define a clone (see Methods). These clones were then used to construct a phylogenetic tree. CCFs of small mutations were also calculated and used to supplement the phylogenetic trees in a subset of cases, refining the inference of subclonal structure. We also compared affected alleles in overlapping CNA from separate tumour regions (described previously as mirrored subclonal allele imbalance, MSAI) which is indicative of parallel evolution^6^.

Overall we observed copy number changes common to WT such as recurrent 1q gain (~28%, frequently observed as subclonal ^10^), 1p loss (~12%), 4q loss (~16%), 8 gain (~22%), 12 gain (~22%), 14q loss (~19%), 16q loss (~17%) and 17p loss (~17%, Figure 1A). Inferred subclone copy number profiles displayed the copy number diversity of WTs (Figure 1B), however many cases contained zero CNA events (14 separate tumours, including 3 non-WTs). Two cases were inferred to be tetraploid (MX-11 and DA-8).

**Figure 1:**
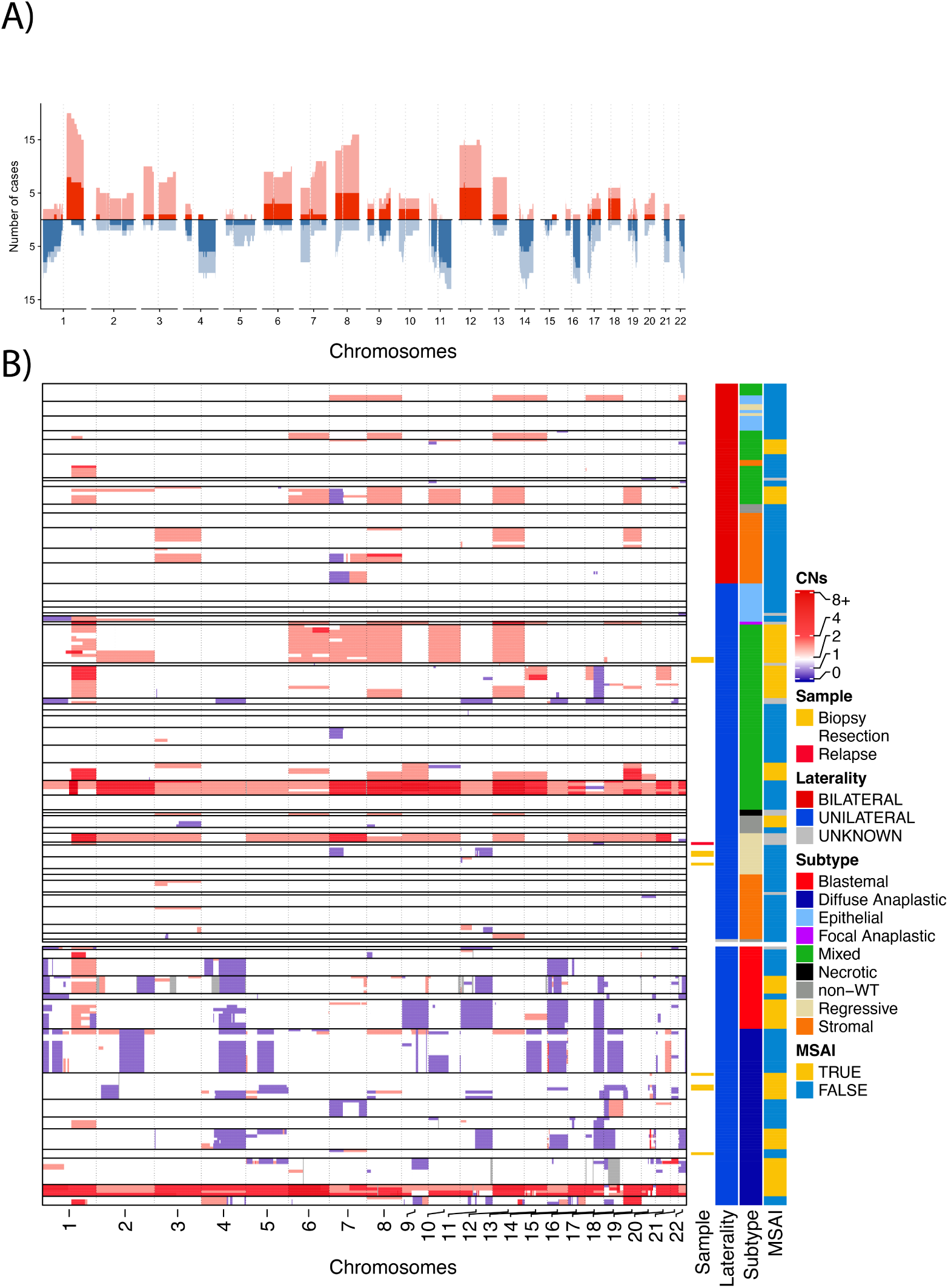
Across 64 paediatric kidney cancers we mapped the prevalence (A) of copy number gains (red) and losses (blue) and the number of cases in which they were clonal (opaque) or subclonal (transparent). Copy number calls across inferred clones are also displayed alongside annotation of sample type, laterality, type of WT (if applicable) and if MSAI is present in the case (B).

### 3.1 Parallel evolution of copy number changes

Strikingly, we observed several examples of parallel evolution in WTs indicating potentially strong selection of CNAs in a variety of contexts. These included parallel evolution between bilateral tumours, independent tumours in the same kidney and within the history of single-origin tumours.

Figure 2A represents a remarkable example of parallel evolution in both a bilateral tumour and between two possibly independent tumours in the same kidney. The patient (BI-6) presented with bilateral WT with an epithelial type histology in both kidneys. In total, 5 samples were taken from the case including 2 from the left kidney and 3 from the right kidney. All samples contained a chromosome 19q uniparental disomy (UPD) event, however it was detected with three unique breakpoints, indicating that the event most likely occurred three times. All samples from the right kidney contained an identical breakpoint, however the breakpoint boundaries in the two samples from the left kidney were unique as compared to the right kidney (supporting independent tumours in different kidneys) but also in respect to each other. This suggests that the left kidney may have contained two tumours, indicating the patient presented with 3 independent tumours, each with the parallel evolution of 19q UPD. This indicates a strong selection for 19q UPD, that each time led to the formation of an epithelial type WT.

**Figure 2:**
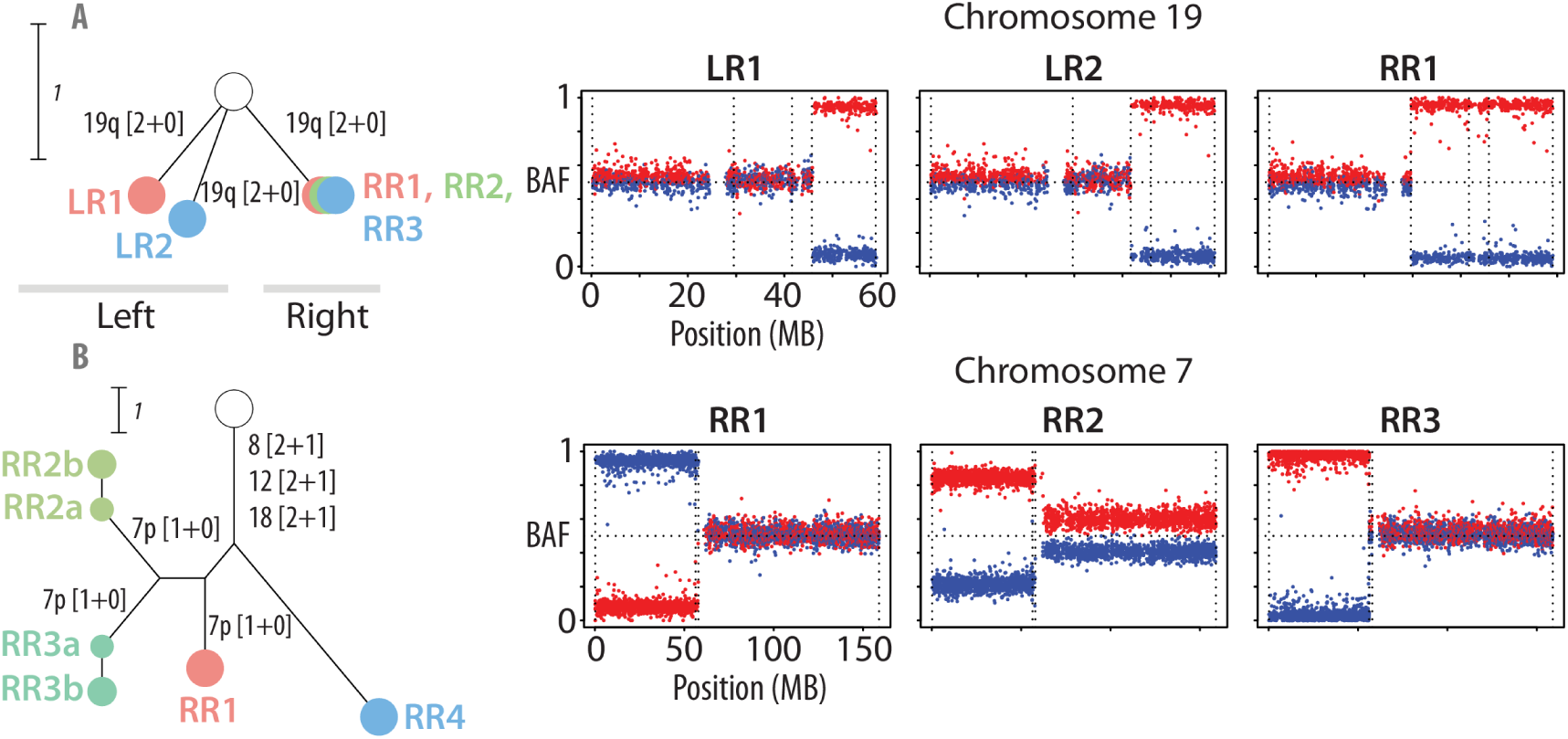
Evidence for parallel evolution in WTs. BI-6 (A) represents a bilateral cancer in which an epithelial type tumour has developed in both kidneys. The case also represents parallel evolution both within two potential tumours within the same mass (left) and between tumours of a different laterality. The only CNA in this case is a 19q UPD event. However, across the five tumour samples taken, three different breakpoints of the chromosome 19q UPD are recorded, one affecting all samples taken from the right (RR1–3) and a unique breakpoint in both samples taken from the left tumour. BI-11 shows parallel evolution within a single tumour (B). Samples RR1–3 all contain unique chromosome 7 loss events. The losses in RR2 and RR3 have different breakpoints and despite the events in RR1 and RR2 affecting the same breakpoint (insofar as is detectable on the SNP array), the event affects different alleles in both samples (as represented by the colour of the phased SNPs).

Another example of parallel evolution is displayed in Figure 2B in a mixed-type WT taken from the right kidney of a patient (BI-11) that presented with bilateral WT. The tumour was sampled 4 times and shows three subclonal 7p loss events. These are supported by the fact that one sample contains a unique break-point boundary (RR3). Two more samples (RR1 and RR2) possess a loss with identical breakpoints, however analysis of MSAI identified the loss as affecting separate chromosomes in these two samples. This indicates there is a strong selective pressure for chromosome 7p loss that is apparently not allele specific.

Three out of seven of the bilaterally sampled cases showed clear parallel evolution between tumours in separate kidneys. In addition to independent 19q UPD events in BI-6 (Figure 2A), the other two cases (BI-4 & BI-7, Supplementary Data 3) showed separate acquisition of 11p UPD in the separate, contralateral tumours, all dominated by stromal differentiation (three stromal tumours and a mixed tumour with 60% stroma – BI-4 right tumour, Supplementary Data 4) implicating 11p UPD as an important initiating event, as previously described ^10^, particularly in tumours dominated by stroma.

We also found six separate instances of parallel evolution in two independent tumours occurring within the same macroscopic mass when assessing CNAs – a phenomenon we observed previously in a single case ^10^. Three of these cases (BI-3, BI-6 and BI-13) were in patients that presented with bilateral tumours, which indicates a strong predisposition to developing multiple lesions both across different kidneys and in the same kidney. The remaining cases were in unilateral WTs (DA-4 and RE-4) and a case of renal cell carcinoma (nW-2).

Furthermore, there were a large number of cases with evidence of parallel evolution of CNAs in a single tumour; in four of these cases, this was parallel evolution of chromosome 1q gain (MX-1, MX-3, BL-4 and BL-6).

### 3.2 *CTNNB1* is under strong selection in a subset of Wilms tumours

*CTNNB1* was mutated in nine of thirty cases that were sequenced using WT-TES. Five of these patients presented with bilateral WT and the majority of the tumours with a *CTNNB1* mutation were stromal type WT (five stromal, three mixed and one blastemal). *CTNNB1* mutations were apparently subclonal in six of nine cases (Supplementary Data 1). In three of these cases multiple *CTNNB1* mutations were present indicating strong and late selection for cells with mutant *CTNNB1*. All three cases exhibiting this parallel evolution of *CTNNB1* mutation were stromal type WT. In ST-5, four separate mutations in *CTNNB1* were discovered in five samples, with each sample containing cells with a mutation in one of the two most common hotspots in *CTNNB1* (amino acids 41 and 45) that prevent phosphorylation of the amino acid residues causing β-catenin stabilisation (Figure 3A–B) ^32^. These *CTNNB1* mutations occurred following a *WT1* frameshift mutation and subsequent 11p uniparental disomy (UPD). Overall this suggests three clones are growing in parallel with four unique *CTNNB1* mutations. Additionally, in BI-7, the right sided stromal tumour contained a K335I mutation in three of the four samples (RR2–RR4), whereas one sample (RR1) contained a S45P mutation (Figure 3C). Additionally, BI-10 displayed parallel evolution of *CTNNB1* as three samples contained a S45Y mutation and one sample contained a S45del in frame deletion (one sample contained no detectable *CTNNB1* mutation). These *CTNNB1* mutations developed in a background of a clonal E1147K *DROSHA* hotspot mutation.

**Figure 3:**
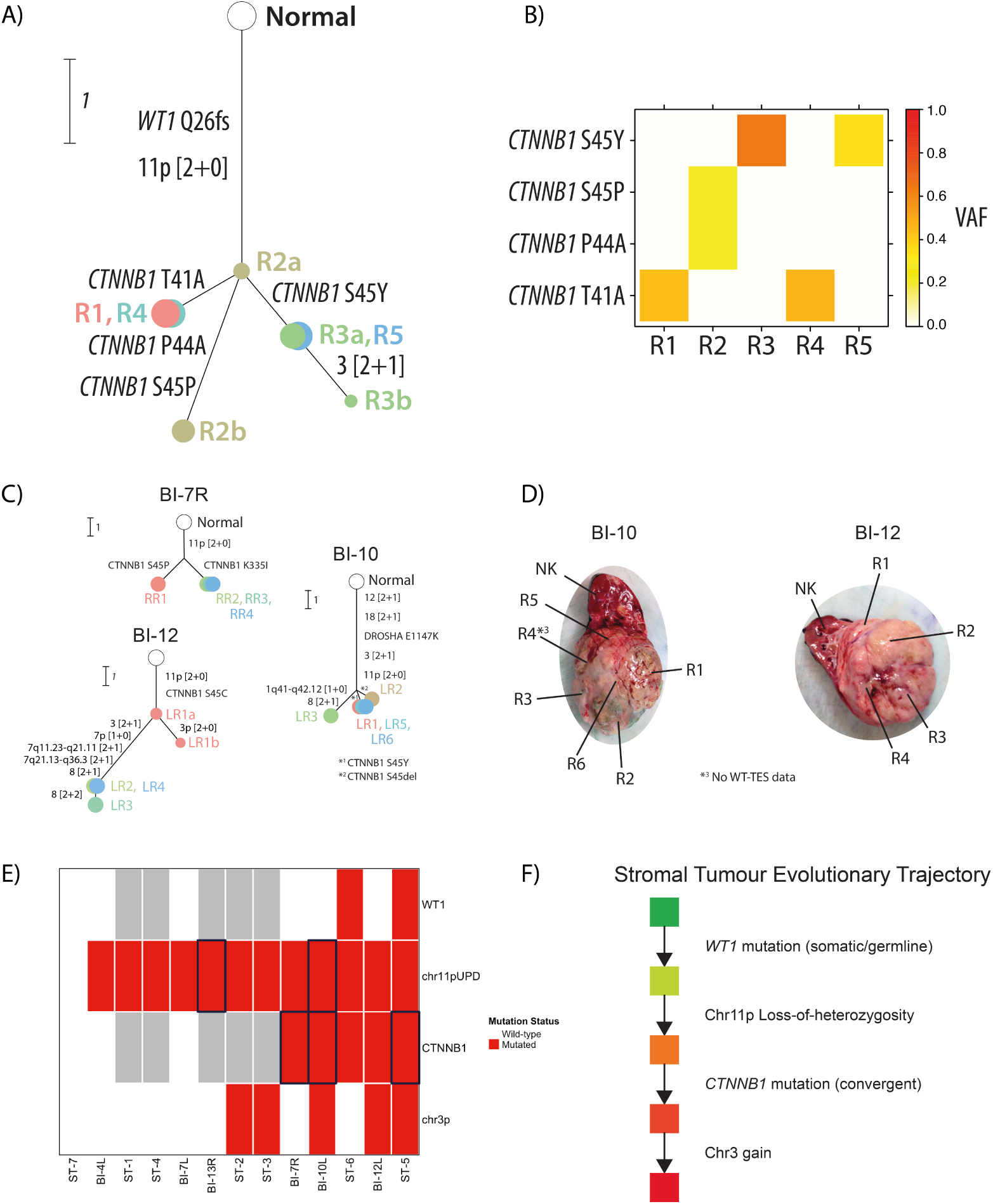
Evidence from the 13 stromal tumours in this study (including six tumours that are one of two bilateral tumours) suggests that stromal tumours undergo a consistent evolutionary trajectory. (A) ST-5 shows parallel evolution of CTNNB1 mutations. Here following a truncating WT1 frameshift mutation and a 11p UPD event, four CTNNB1 have been acquired, as observed in five separate tumour pieces. The heatmap (B) shows the recorded VAF of the CTNNB1 mutations. (C) The evolution of stromal WT frequently involves WT1 mutation, 11p UPD and parallel evolution of CTNNB1 mutation and 3p copy number changes. (D) Inferred clones can be mapped to the locations on the tumour from which they originate. (E) Stromal WT evolution is supported by the presence of several key events in our cohort that also evolve in parallel (black boxes, grey = data not available). (F) The evolution of a typical stromal WT begins with an initial WT1 mutation, followed by a LOH event that affects both the WT1 and IGF2 loci. Following this there is an acquisition of CTNNB1 mutations, sometimes in clones evolving in parallel, indicating late but strong positive selection for WNT signalling dysregulation in stromal tumours. A subset of tumours then proceeded to acquire CNAs in chromosome 3, the majority of which affected the CTNNB1 locus.

### 3.3 A model for the evolution of stromal tumours

Looking more broadly at the context of *CTNNB1* parallel evolution, we observed that the most striking example of a consistent evolutionary pattern were the alterations present in stromal WT. When considering bilateral tumours as individual tumours, all tumours bar one unilateral stromal tumour (12/13, Supplementary Data 3) displayed a 11p UPD event. Chromosome 11p UPD was clonal in all but two bilateral tumours, however the event emerged multiple times through convergent evolution (2/12). In all cases with somatic *WT1* mutation (3/30) the mutation appeared clonal and occurred in tumours that also had *CTNNB1* mutation, two of these cases were stromal tumours (the remaining 6 sequenced stromal tumours did not contain a somatic *WT1* mutation). Five sequenced stromal tumours contained a *CTNNB1* mutation (5/8, Figure 3E). Stromal histologies were exclusively affected by convergent evolution of *CTNNB1* mutation (n=3), indicating *CTNNB1* mutation as a late event in stromal tumour evolution that is under strong selection. In the remaining two stromal tumours with *CTNNB1* mutation, the mutation appeared clonal. Many 11p UPD events were followed by 3p changes in copy number analysis, which may suggest selection due to the acquisition of a *CTNNB1* mutation, as supported by the fact CNA affecting the *CTNNB1* locus appeared exclusively in the *CTNNB1* mutated tumours in the sequenced cohort. This data indicates the observation of progressive alterations in stromal WTs (Figure 3F). A typical evolution of a stromal tumour likely involves *WT1* mutation followed by an 11p UPD event that involves 11p13 (therefore the *WT1* locus, as well as *IGF2* on 11p15). This then leads to strong selection for *CTNNB1* mutation as well as chromosome 3 gains or UPD events (no chromosome 3 losses were observed in stromal WT that had a 11p UPD event).

### 3.4 Diversification in Wilms tumour evolution

Multiregion WT-TES revealed a *TP53* mutation in each of the seven assayed diffuse anaplastic tumours. In four cases (DA-2, DA-4, DA-7, DA-9, Supplementary Data 1) *TP53* mutation was heterogeneous and was associated with strong diversification. In three cases displaying subclonal *TP53* mutational status, the mutation was associated with the accumulation of several extra CNA events, acquired after the CNAs found in other regions (Supplementary Data 3). This suggests the development of subclonal genomic instability. The mutations causing this increase in subclonal CNAs were E285K (R3, DA-2), P82fs insertion (R1, DA-7) and H179L (R2, DA-9). In two cases (DA-2 and DA-9) these occurred following only a 19q UPD event. In two additional regions of DA-7 (R2 and R4, total 4 regions sequenced), we also detected two more unique *TP53* mutations, R342* (R2) and G245S (R4), showing striking parallel evolution. These regions did not exhibit large CNA accumulation as in R1, suggesting that either only particular *TP53* mutations cause this effect, that additional unknown changes are also required, or that *TP53* mutant clones are under positive selection prior to the accumulation of CNAs. Each region containing a subclonal *TP53* mutation also exhibited LOH at the locus in all three cases.

Heterogeneity in *TP53* status was related to phenotypic difference in DA-4 (Figure 4). Here R3 displayed an anaplastic phenotype whereas R1 and R2 contained mostly blastemal cells, therefore the tumour displayed two high risk WT phenotypes in the same mass. From the perspective of CNAs these phenotypically distinct regions share no changes (losses in 15q and 16q have different breakpoint boundaries). However, WT-TES revealed a *DROSHA* E993K mutation in all samples indicating that the regions were related and not independent. *DROSHA* E993K mutation has been previously reported in WT^33^. The anaplastic region (R3) uniquely contained a *TP53* E339del mutation and the blastemal regions possessed a *SIX1* Q117R mutation, a hotspot previously associated with blastemal tumours ^17^. This phenotypic disparity is likely a result of large mutational divergence following the *DROSHA* E993K mutation – which may have been a field effect causing mutation. This case represents the large potential for diversity in WT with different regions strikingly phenotypically distinct, driven by almost independent yet related genotypes. Furthermore, the normal kidney sample used as reference in DA-4 shows 7/571 reads reporting the clonal E993K mutation in *DROSHA*, there is not a single read reporting the other non-clonal point mutations discovered in *DROSHA*, *SIX1* and *GNAS* in the normal kidney sample. Ergo, 2.45% (1.19% – 5.02%, binomial confidence interval, Wilson method) of the cells in this normal kidney sample may already have the E993K *DROSHA* mutation. Hence, the kidney may be mosaic for the *DROSHA* mutation that then initiates two nearly independent tumours.

**Figure 4:**
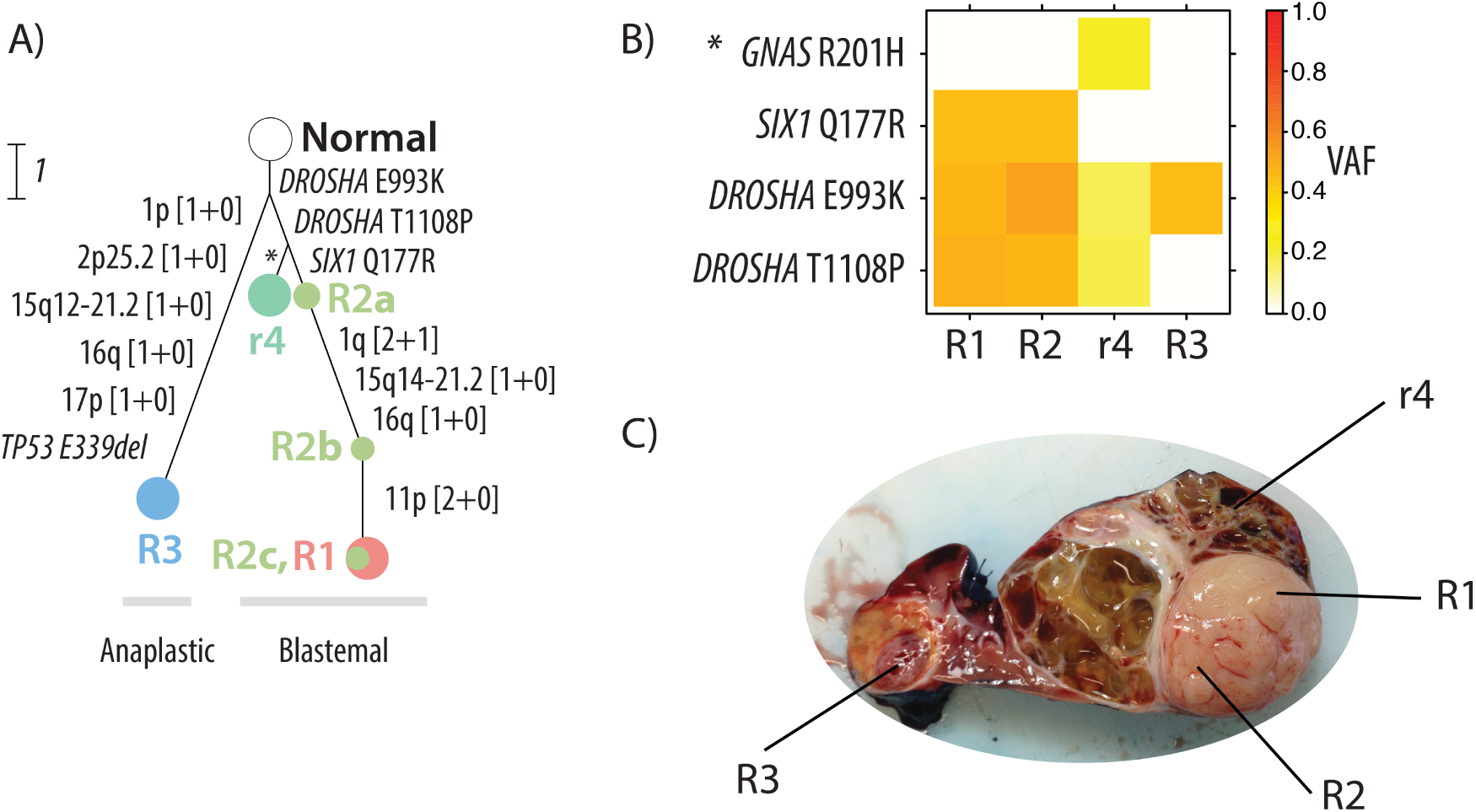
Strong early diversification in a WT case (DA-4). Tumour regions R1–2, r4 follow a separate evolutionary trajectory to R3 following the acquisition of a clonal DROSHA E993K mutation (A). R1–2, r4 acquire an additional DROSHA mutation (T1108P) and R1, R2 further diversify acquiring a SIX1 Q177R mutation and CNAs. R1 and R2 showed blastemal histology. R3 developed unique CNAs (including 17p loss) and a TP53 E339del mutation. R3 showed diffuse anaplasia and led to the case being diagnosed as such. The heatmap (B) shows the VAF of each point mutation across the regions supporting this phylogeny. The photograph of the tumour (C) shows that the regions are located in distinct sites. R1 and R2 form part of an outgrowth. R3 is spatially separate from R1–2, r4. SNP array data was not available for r4.

BI-11, a mixed type WT taken from a patient with bilateral disease, also displayed phenotypic diversity associated with underlying genetic heterogeneity. Here RR4, exclusively contains a N387K *CTNNB1* mutation and the region presented with 95% stromal cells. The other tumour pieces RR1–RR3 share a R358* *AMER1* mutation, present in only 2% of reads in RR4. Regions RR1–RR3 contained a mixed-type histology. These tumours are related by shared whole chromosomal gains of 8, 12 and 18, yet show divergence both genetically and phenotypically.

### 3.5 UPD of chromosome 19q is a potential link between epithelial and diffuse anaplastic tumours

We observed LOH affecting the telomeric end of chromosome 19q in 15 cases and in 11 cases the LOH was clonal. Of the cases in which the chromosome 19q LOH event was clonal, eight of these events were UPD [2+0], the other three were losses. Interestingly, of the eight cases in which clonal UPD of 19q was observed, four were epithelial type WT (BI-6, EP-1, EP-2, EP-3), three were diffuse anaplastic type (DA-2, DA-6, DA-7) and one was a mixed histology case (MX-10). Additionally, two of these three diffuse anaplastic cases were dominated by epithelial histology (DA-6 – 70%, DA-7 – 80%).

Remarkably, in all four epithelial tumours containing a 19q UPD event, it was clonal and the only observed change (Figure 5A, as highlighted previously, BI-6 developed three tumours each only with a chromosome 19q UPD event). Thus, from a CNA perspective, 19q UPD appears sufficient to form an epithelial WT.

**Figure 5:**
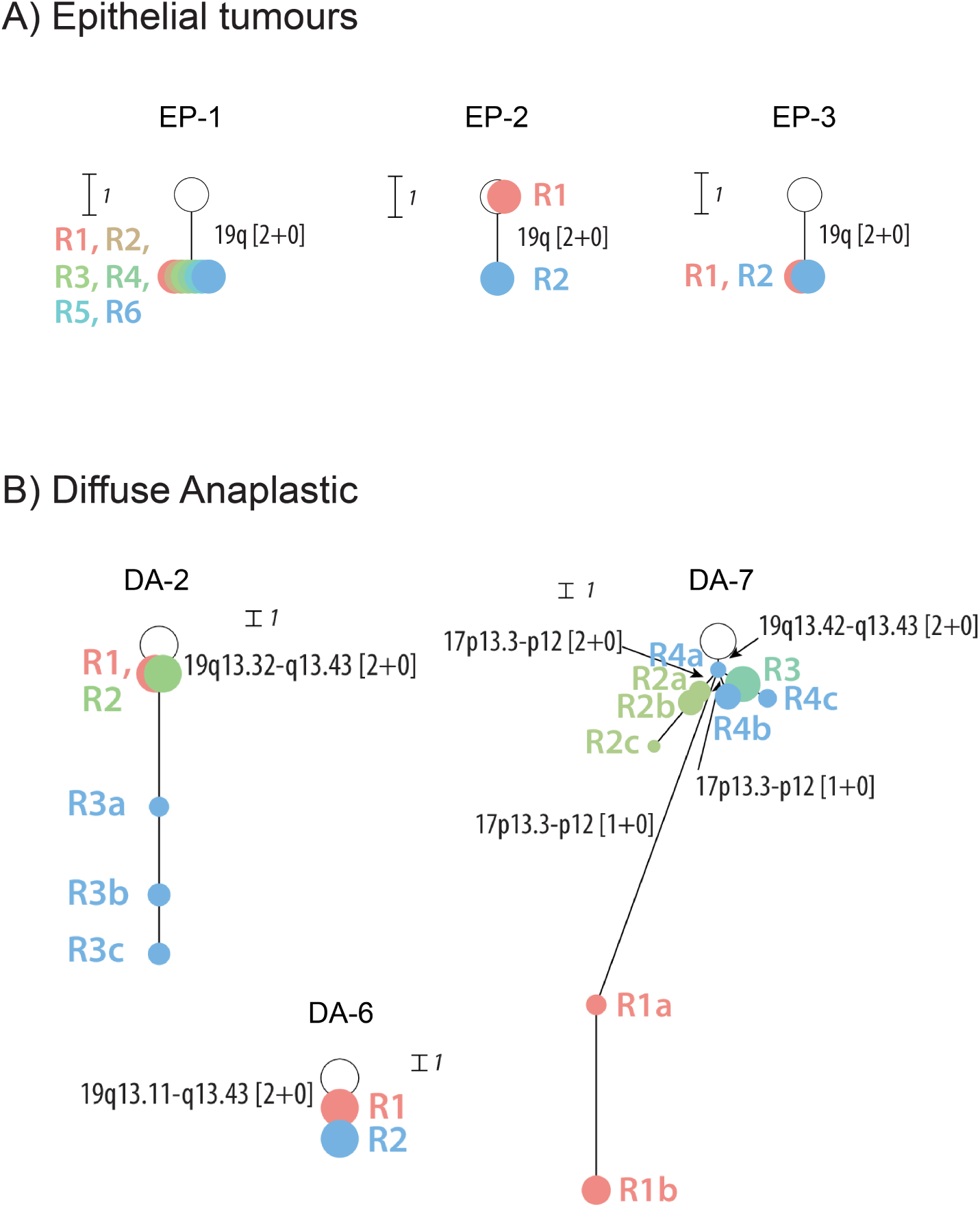
A subset of epithelial tumours (A) and diffuse anaplastic tumours (B) shared the common feature of early 19q UPD. All tumours had observed ancestral clones containing only 19q UPD.

Additionally, in all three diffuse anaplastic tumours with a clonal 19q UPD event there is at least one tumour sample in each in which the aberration is the only CNA observed (DA-2, DA-6) or the only observed CNA which is completely clonal (DA-7 R4a, Figure 5B) proving the CNA was an initiating event. We hypothesise that these diffuse anaplastic cases may be further evolved epithelial tumours.

By examining the 19q UPD events in each tumour in which they are clonal, the genomic location which is affected can be mapped. In all eight cases the UPD event affects the entirety of chromosome 19q13.43. All chromosome 19q13.43 clonal UPD events are not affected by MSAI, indicating that the event is allele-specific (Figure S1).

### 3.6 Total number of evolutionary copy number events across multiple regions is linked to event free survival in high risk WTs

We obtained follow up data for 53 cases in our cohort and we were able to test if general features of the evolution of these cancers were prognostic. We assessed both total number of evolutionary CNA events (ECEs), a proxy of genomic instability, across the series as well as percentage of subclonal ECEs, as possible predictors of relapse/death. For both predictors we generated a ROC curve to test for the relationship between specificity and sensitivity of these predictors for event-free survival. Moreover, the ROC curve allowed us to choose a threshold for each predictor that seeks to maximise sensitivity and specificity (Figure S2).

ROC analysis of ECEs produced a sensitivity/specificity with the greatest distance from what is expected randomly at 11 events (AUC = 0.65, Figure S2A). Using this value to differentiate between patients in a category of high/low number of ECEs significantly predicted event-free survival (*p* = 0.00014, log-rank test, Figure S2B). This indicates that genomic instability is predictive of event-free survival in paediatric kidney cancer. ROC analysis indicated ECE subclonality was less predictive (AUC = 0.57, Figure S2C).

To explore ECEs and prognosis further, we assessed the ECEs across different histological risk types. Generally, we observed that high risk tumours (blastemal and diffuse anaplastic) have a larger number of ECEs (median number of events = 29, range 4–110, Figure 6A). The median numbers of ECEs for intermediate and low risk were 4 and 3 events, respectively, and the median number of ECEs in a small set of non-WT paediatric kidney cancers was 0. The range of ECEs was large for intermediate type WTs (0–71) and six tumours, shown as outliers in Figure 6A, have more than 15 ECEs.

**Figure 6:**
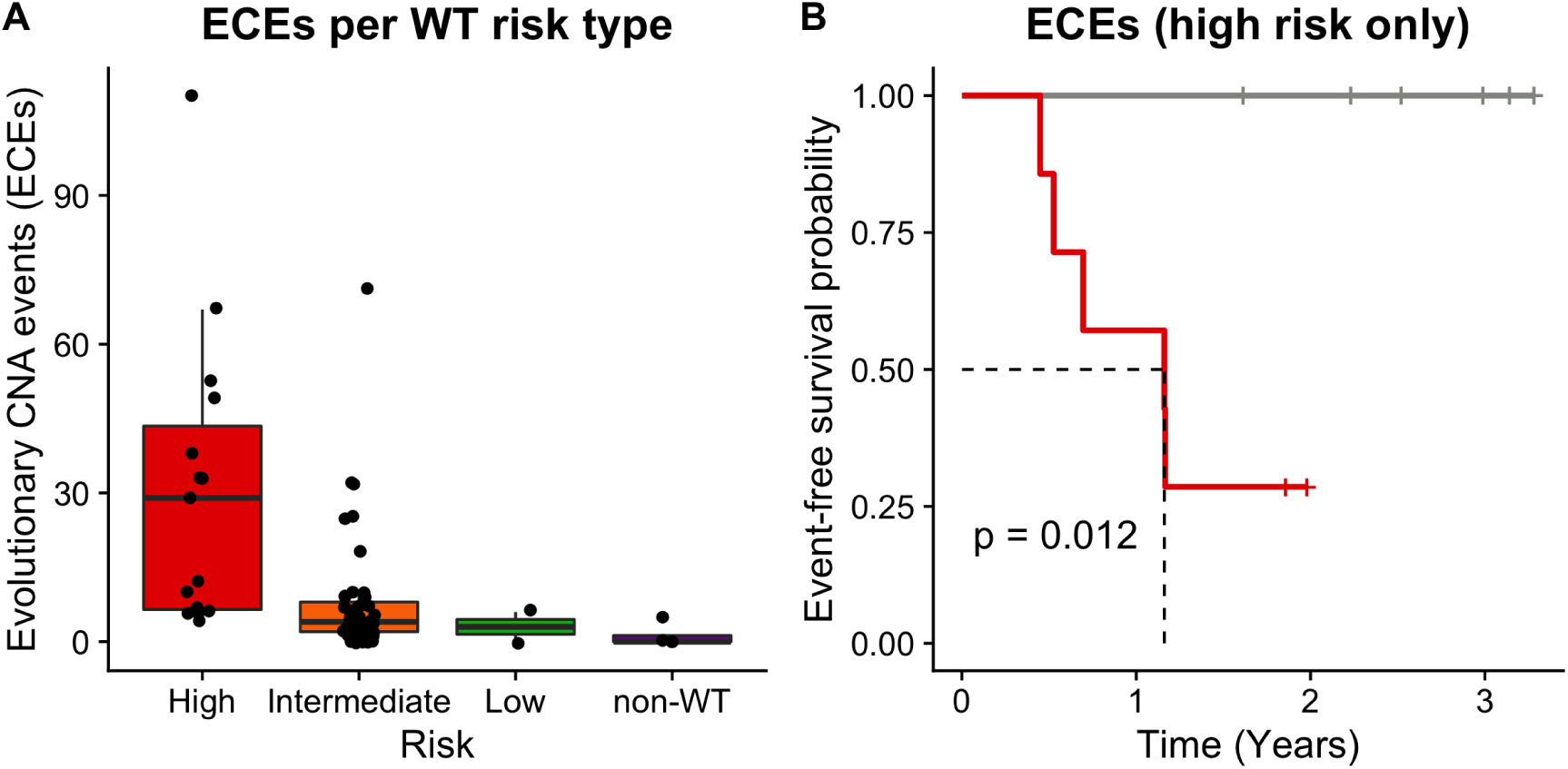
The number of ECEs per histological WT risk type is variable (A). Low risk WTs and other paediatric kidney cancers have very few ECEs over the course of their evolutionary history (low risk median ECEs = 3, n = 2). In stark contrast, high risk WTs show many ECEs, with a median number of ECEs of 29 and a range of 4–110 (n = 15). Intermediate risk WTs generally have much fewer ECEs than high risk WTs (median = 4, n = 43), however there are outlier cases that have more than 15 ECEs. Event free survival analysis of high-risk cases only, split by high (red) or low (grey) number of ECEs (threshold of 11 calculated by ROC analysis, Fig. S2) showed that high-risk tumours with a high level of genomic instability had worse event free survival than high-risk tumours with low genomic instability (B, log-rank test, p = 0.012, n = 13).

We then tested if ECEs could be prognostic within high risk WTs themselves. Using a threshold of greater than 11 ECEs (as calculated on the whole cohort using ROC analysis) to classify cases as high/low instability we performed a log-rank test on the Kaplan-Meier curves of the high risk cases. Total number of ECEs in these cases appeared to be prognostic of relapse/death (*p* = 0.012, log-rank test, Figure 6B). This suggests that the number of ECEs in the history of high risk tumours can be used to distinguish between cases that are likely to relapse or not, adding further power to risk stratification (n = 13).

## 4 Discussion

Here we present a large cohort of WTs and other paediatric kidney cancers, in which the majority of patients are multi-sampled, alongside the detailed characterisation of their evolutionary progression and its association with histopathological and clinical data. These results support and build on recent publications of similar cohorts of WT patients ^10–13,34^ and show that the evolutionary patterns of WT are frequently consistent and robust, especially in relation to the histopathological types and risk groups.

We quantified the number of observed CNA events across all WT risk categories and were able to show the prognostic potential of these gene dosage modifications. This supports previous observations that mean branch length is predictive of survival across a cohort of neuroblastomas, WTs and rhabdomyosarcomas^13^ as well as the observation that an imbalance burden of *>*100Mb is prognostic in WTs ^11^. These data support the concept that evolvability may be a key feature of high risk WTs (and potentially paediatric cancers in general) and observing a large number of CNA events across multiple regions may provide evidence that the tumour has explored a significant amount of its fitness landscape and is capable of adapting to changing fitness pressures. It may be more important to measure evolvability using CNAs as WTs have the potential to exhibit a large burden of chromosomal alterations but rarely present with large burdens of point mutations and indels ^17^. Additionally, we show that this feature may be able to differentiate patients with a greater risk of relapse within the high risk category itself and may allow for alternate treatment regimes that are less cytotoxic in patients with high risk WT that exhibit low evolvability. We argue that a shift in conceptualising evolutionary features away from merely attempting to describe ‘heterogeneity’, particularly of individual biomarkers, provides clearer theory behind clinically useful features; indeed percentage of subclonal CNA events was a less useful feature in our study for differentiating relapsing patients. Overall, a larger study of high risk types is required to prove the usefulness of assessing the acquisition of ECEs across multiple regions. Measurements of total chromosomal instability will likely supersede single CNA biomarkers, such as 1q gain, in prognostic usefulness as chromosomal instability measures a fundamental characteristic of the cancer. Indeed the prognostic potential of 1q gain ^35^ may be a proxy of increased chromosomal instability.

The evolutionary patterns of WT types are a manifestation of their adaptability as well as their underlying biology. It is clear that diffuse anaplastic WTs are defined by large acquisitions of CNAs following late, subclonal *TP53* mutation and that this may happen in parallel several times in the population and not be clonal at surgery, as others have also shown ^12–14,36^. The fact that diffuse anaplastic tumour populations may still contain cells with few CNAs and no *TP53* mutation indicates that these events do not provide proliferative advantages large enough to outcompete the ancestral genotype completely, yet may be linked to increased chance of relapse due to the untapping of new adaptable potential. This adaptability may also be acquired by blastemal tumours, however it does not appear to be driven by *TP53* mutation. Furthermore, in diffuse anaplastic tumours *TP53* mutation did not always necessitate a large expansion in CNA events (e.g. DA-4 and DA-7), indicating that other conditions may need to be met, that only certain *TP53* mutations produce this change, or that the acquisition of large number of CNA events is produced by a rare event.

In this study we frequently found clonal chromosome 19q UPD in both epithelial and diffuse anaplastic samples. These events were likely to be allele-specific as they never displayed MSAI. The fact that several cases of diffuse anaplastic WT presented with clones with only 19q UPD is strong evidence that it is an early event in these tumours, relative to additional CNAs, as well as being recurrent in epithelial WTs. The late acquisition of *TP53* mutation in diffuse anaplastic tumours and the strong association to the presentation of anaplastic cells indicates that a tumour observed earlier in its evolutionary trajectory, prior to *TP53* mutation, may have been classified as a different histological type. The evolutionary link to chromosome 19q UPD indicates that epithelial tumours may have a higher chance of producing diffuse anaplasia. The importance of chromosome 19q UPD in the early stages of epithelial tumour is strongly exemplified by the parallel acquisition of three independent 19q UPD events in a case of bilateral epithelial WTs, as well being the sole CNA event in 3 of 5 unilateral epithelial tumours. Anaplasia and *TP53* mutation has been previously reported in epithelial tumours with *TRIM28* mutation, located on chromosome 19q supporting the hypothesis that *TRIM28* could be associated with both histologies ^37–39^. This may indicate that some *TRIM28* mutant tumours with intermediate risk may at some point transition to anaplasia.

The acquisition of CNA events that evolve in parallel indicate strong selective pressures ^6^ and therefore provide the clearest picture of the underlying biology of tumorigenesis in WTs. In stromal WTs we observe parallel evolution of chromosome 11p UPD and in particular *CTNNB1* mutation, supporting observations seen in two WT patients with germline *WT1* mutation ^34^. Stromal WTs show a clear trajectory from *WT1* mutation to 11p UPD and subsequent *CTNNB1* mutation and 3p CNAs. There is likely to be strong epistasis at each point in this trajectory for the subsequent alteration that represents consistent changes in the fitness landscape and successful exploration to produce a stromal phenotype. The presence of this succession of events also occurred in a mixed type tumour (BI-11) and produced a region of the tumour rich in stromal content, indicating that cells may explore this pathway alongside other non-stromal type WT cells. Similarly, we observed an example of putative diffuse anaplastic and blastemal trajectories evolving in parallel in a single tumour mass, indicating that parallel trajectory exploration may be common. The consistency of the stromal WT evolutionary trajectory provides many potential opportunities. For example, it may be used to stage the progression stromal WTs. Additionally, the convergent evolution of *CTNNB1* mutation may allow for the quantification of selection of many different *CTNNB1* mutants in the same individual. Revealing the *CTNNB1* mutations that provide the greatest fitness advantages. This could help determine the prognostic value of different *CTNNB1* mutants and help to predict the clone that may present at recurrence ^40^.

The fact that key mutations are acquired in parallel, late in the evolution of different WT types is highly informative of the tumorigenesis of the cancer and is even a predictable feature of WTs given their large size ^41^. The consistency of these evolutionary trajectories highlights the benefit of characterising WT evolution and allows us to measure inherent characteristics of the WT cancer population that will move us beyond single genetic biomarkers. Furthermore, highlighting how early alterations to transcriptional programmes/cell signaling appears to produce a selective advantage for mutations that alter other cancer-related signaling pathways exposes the sequential and repeatable biological rewiring that occurs in WTs.

## 5 Conclusions

Here we provide an in-depth evolutionary analysis of a large cohort of WT patients with reference to histological subtypes and clinical outcomes. We show that parallel evolution highlights consistent evolutionary patterns within subtypes. In addition to characterising subtypes, evolutionary analysis also provides links between subtypes, as exemplified by 19q UPD in epithelial and diffuse anaplastic WT. This characterisation of evolution across diverse WT clinical presentation allows us to fundamentally understand these subtypes and moulds an expectation of their evolutionary progression. The fact that multiple pathways of evolution can be observed within the same tumour mass explains why different histological phenotypes can be detected within the same patient and indicat that evolutionary pathways can diverge early in tumour evolution. Finally, by using our evolutionarily informed analysis of CIN for prognostication we are able to show that WT patients with a large number of ECEs have poor outcomes and that this relationship holds in high-risk patients. We suggest an evolutionarily informed measurement of CIN (i.e. ECE quantification) should be considered as an approach to identify the most aggressive high-risk WTs, as opposed to single CNA biomarkers.

## Abbreviations

AUC: Area under the curve
BAF: B-allele frequency
CCF: Cancer cell fraction
CCLG: UK Children’s Cancer and Leukaemia Group
CNA: Copy number alteration
DHPN: Diffuse Hyperplastic Perilobar Nephroblastomatosis
ECE: Evolutionary copy number alteration events
EM: Expectation maximisation
H&E: Haematoxylin and eosin
IMPORT: Improving Population Outcomes for Renal Tumours of Childhood
LOH: Loss-of-heterozygosity
LRR: Log2 R ratio
mBAF: Mirrored B-allele frequency
MSAI: Mirrored subclonal allelic imbalance
ROC: Receiver operating characteristic
TES: Targeted exon sequencing
UPD: Uniparental disomy
VAF: Variant allele frequency
WT: Wilms tumour

## Declarations

### Ethics approval and consent to participate

The research was approved by the NHS National Research Ethics Service Committee London - London Bridge (12/LO/0101). Informed consent to participate was obtained from parents or legal guardians of all of participants in the study.

### Publication Consent

Not applicable.

### Contributions

G.D.C., T.D.T., B.M. and W.M. analysed the data and interpreted the results. T.C., R.A.S., R.W., T.D.T., and G.M. generated the data. G.V. and W.M. provided histopathological assessment. G.D.C. and W.M. prepared the manuscript. G.D.C., R.W., N.M.L., K.P.J. and W.M. helped design the study. B.M., T.D.T., N.M.L., K.P.J. and W.M. supervised the study and contributed to the manuscript.

## Acknowledgements

We acknowledge UCL Genomics for their help generating genomic data for this study.

## Supplementary Data

**Supplementary Data 1:** Mutations called on the cohort of sequenced samples.

**Supplementary Data 2:** Copy number alterations and their estimated cellularity.

**Supplementary Data 3:** Phylogenetic trees calculated from copy number alteration data.

**Supplementary Data 4:** Summary of clinical and genomic features per case.

**Supplementary Data 5:** Follow-up information for all cases.

**Supplementary Data 6:** Locations of detected mirrored subclonal allelic imbalance.

---

**Figure S1:**
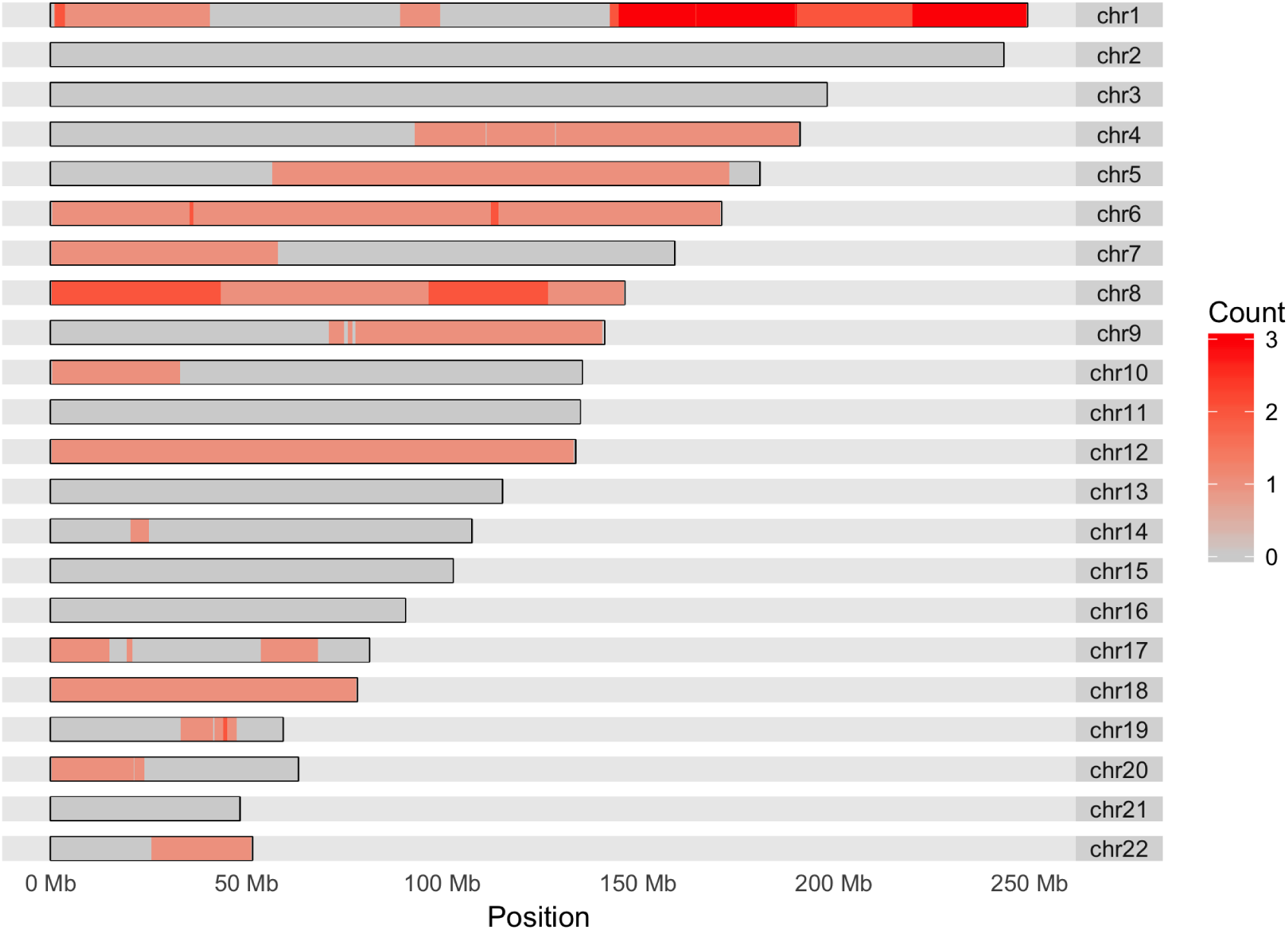
Mirrored subclonal allelic imbalance (MSAI) was detected in 12 cases in our series. It was sparsely distributed across the genome. The most common location for MSAI was in chromosome 1q. Six of the twelve cases with detected MSAI showed MSAI in chromosome 1. Chromosome 11 is not affected by any MSAI events. This indicates that intratumour heterogeneity of chromosome 1q events are not allele specific.

**Figure S2:**
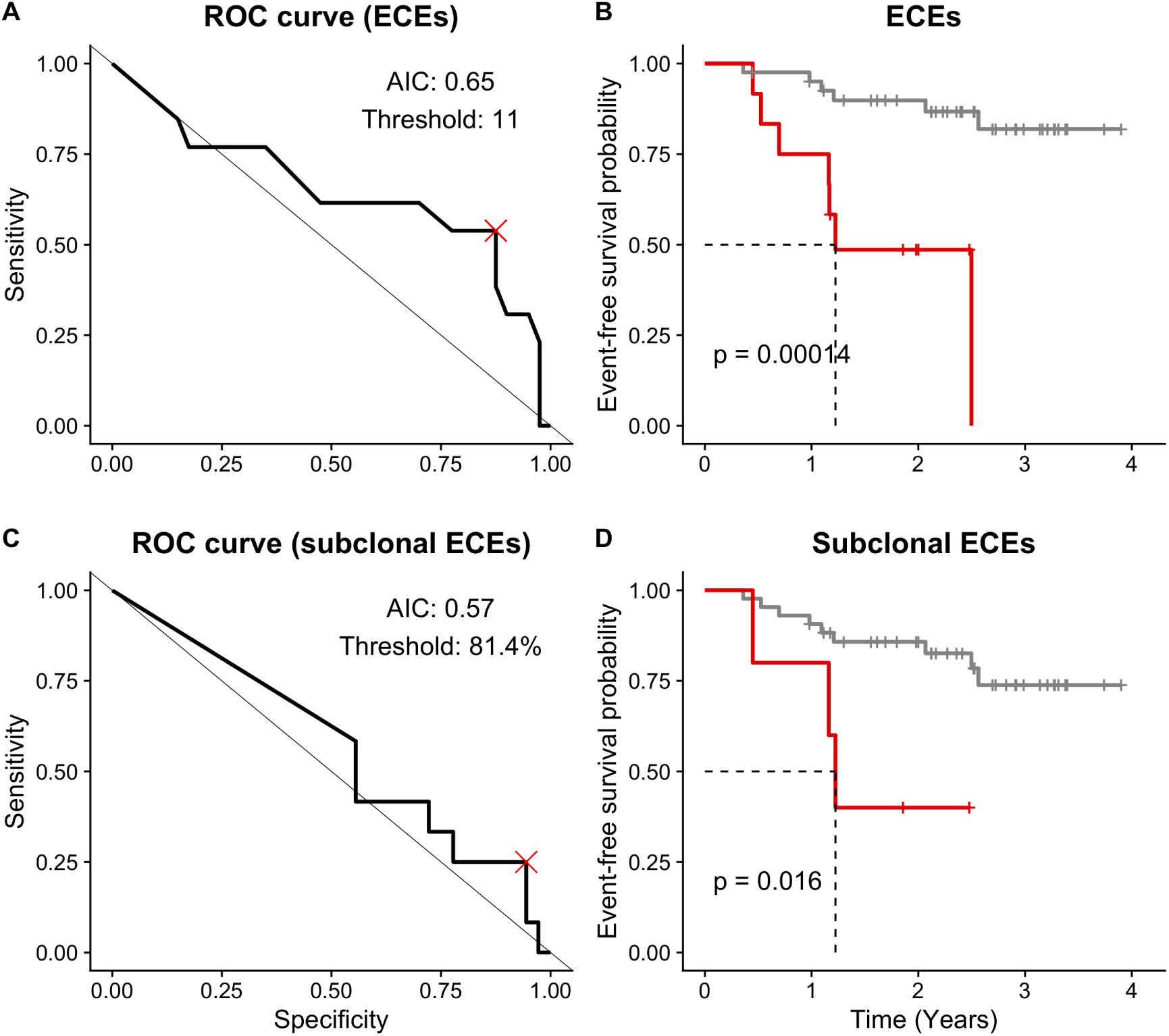
Genomic instability, measured as total number of evolutionary CNA events (ECEs) in tumour evolution, and percentage of CNA which are subclonal, appear to be prognostic for event-free survival. To investigate this both the number of ECEs per tumour and fraction of subclonal ECEs were used to perform ROC analysis (A, C), here the number of ECEs has a greater area under the curve (AUC) than fraction subclonal (0.65 v 0.57) indicating that total ECEs is more predictive of event-free survival than subclonal ECEs. Based on the ROC analysis the threshold that produced the value furthest from the line representing a random predictor (x = 1 - y) in ROC space was taken (as shown by the red cross). This value was 11 events, for total ECEs, and 81.4% subclonal ECEs, for subclonality. Performing a log-rank test on the Kaplan-Meier curves for each predictor (B, D) showed both the total number of ECEs (p = 0.00014) and the fraction of subclonal ECEs (p = 0.016) to be predictive of event-free survival.

**Table S1:**
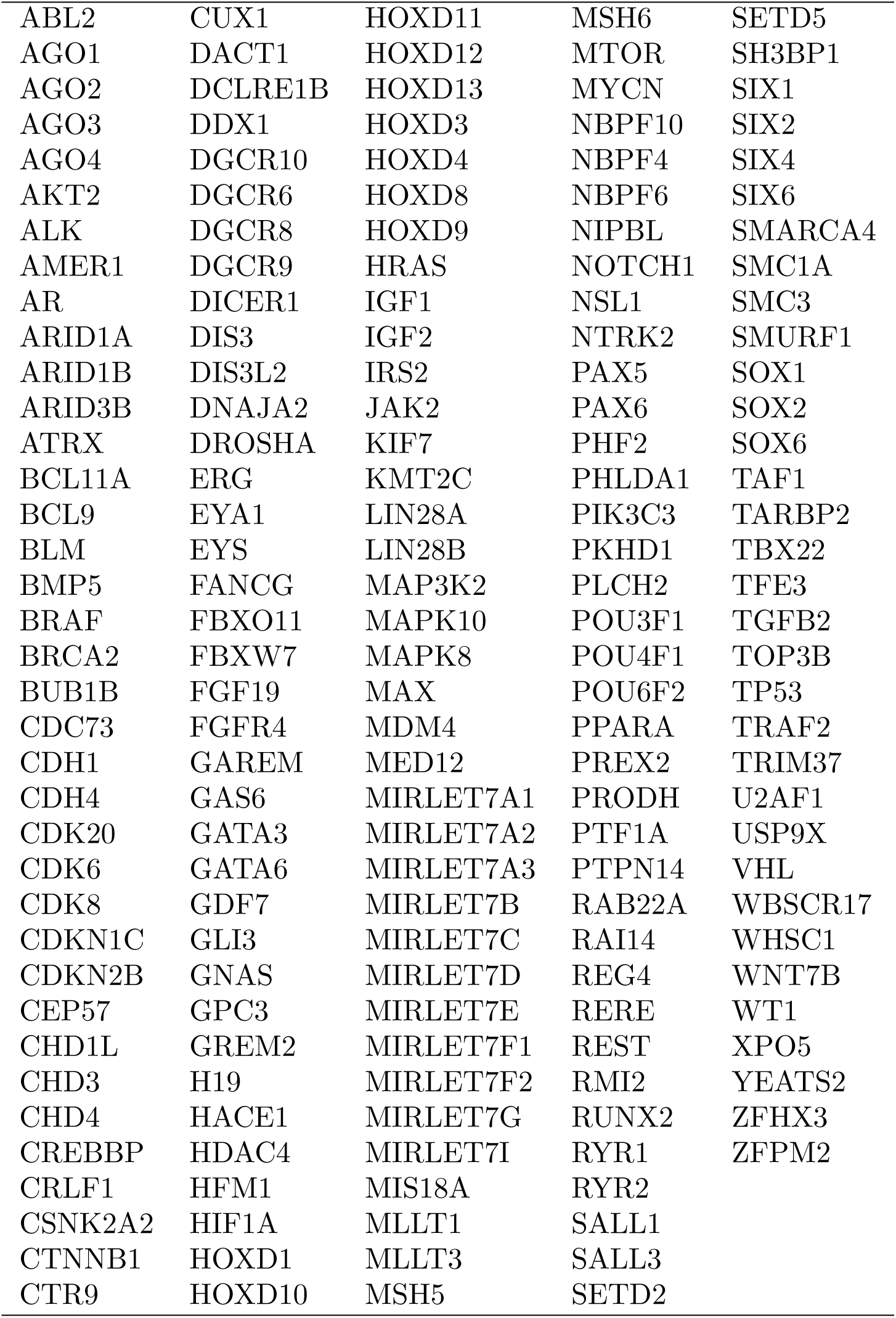
Genes included on the WT-specific targeted sequencing panel (WT-TES).

